## Supplemental Tables for "Teratogenic Drug Topiramate Upregulates TGFβ1 and SOX9 Expression in Primary Palatal Mesenchyme Cells"

### Table S1: List of all 57 proteins with altered expression in Topiramate-treated HEPM cells

*|FC| ≥ 1.15

| *Protein Name* | *Symbol* | *UniProtKB ID* | *Antibody Array ID* | *Fold Change* |
|---|---|---|---|---|
| Checkpoint kinase 2 | CHEK2 | O96017 | 659 | 1.96 |
| Low-density lipoprotein receptor class A domain-containing protein 1 | LDLRAD1 / LRP1 | Q5T700 | 109 | 1.64 |
| Aldehyde dehydrogenase 7 | ALDH3B1 | P43353 | 92 | 1.46 |
| NADPH oxidase 5 | NOX5 | Q96PH1 | 282 | 1.46 |
| Ret Proto-oncogene | RET | P07949 | 490 | 1.42 |
| Cardiac troponin I | TNNI3(cTnl) | P19429 | 146 | 1.39 |
| Transforming Growth Factor beta 1 | TGFB1 | P01137 | 377 | 1.39 |
| Unconventional myosin Id | MYO1D | O94832 | 276 | 1.37 |
| Tyrosine-protein kinase receptor 3 | TYRO3 | Q06418 | 478 | 1.33 |
| Anaphase Promoting Complex Subunit 1 | ANAPC1 | Q9H1A4 | 1093 | 1.32 |
| Superoxide dismutase 1 | SOD1 | P00441 | 996 | 1.31 |
| Oral cancer-overexpressed protein 1 | ORAV1 / ORAOV1 | Q8WV07 | 1078 | 1.30 |
| Growth arrest and DNA damage-inducible proteins-interacting protein 1 | GADD45GIP1 | Q8TAE8 | 1285 | 1.27 |
| Phosphatidylinositol-glycan biosynthesis class H protein | PIGH | Q14442 | 453 | 1.25 |
| CD30 Ligand | CD153 | P32971 | 220 | 1.24 |
| Patched | PTCH1 | Q13635 | 35 | 1.23 |
| Cadherin-10, T2-cadherin | CDH10 | Q9Y6N8 | 415 | 1.21 |
| ATP synthase subunit delta, mitochondrial | ATP5D | P30049 | 261 | 1.19 |
| TNF Receptor1-associated DEATH domain protein | TRADD /TNFR1 | Q15628 | 716 | 1.19 |
| Alcohol dehydrogenase class 4 mu/sigma chain | ADH7 | P40394 | 91 | 1.19 |
| Interferon Regulatory Factor 4 | IRF4 | Q15306 | 730 | 1.18 |
| Mitochondrial 2-oxodicarboxylate carrier | SLC25A21 | Q9BQT8 | 1005 | 1.14 |
| POTE Ankyrin Domain Family, Member G | POTEG (A26C2/3) | Q6S5H5 | 623 | 1.12 |
| Apoptosis regulator BAK / BCL2-Antagonist/Killer 1 | BAK1 | Q16611 | 10 | 1.11 |
| Heat Shock Protein 90-alpha | HSP90A | P07900 | 867 | 1.11 |
| Caspase 10 | CASP10 | Q92851 | 13 | 1.09 |
| Coagulation factor XII (heavy chain,Cleaved-Arg372) | FA12 | P00748 | 1145 | 1.08 |
| Matrix Metalloproteinase-8 | MMP-8 | P22894 | 1210 | 1.05 |
| Abhydrolase Domain Containing 12 | ABHD12 | Q8N2K0 | 83 | 1.03 |
| Galectin 7 | LEG7 / LGALS7 | P47929 | 1261 | -1.05 |
| Cytochrome P450 2W1 | CYP2W1 | Q8TAV3 | 927 | -1.05 |
| Ephrin type-A receptor 6 | EPHA6 | Q9UF33 | 635 | -1.08 |
| Claudin 11 | CLDN11 | O75508 | 181 | -1.08 |
| USP6 N-Terminal Like | USP6NL | Q92738 | 292 | -1.08 |
| Histone Deacetyalase 1 | HDAC1 | Q13547 | 855 | -1.12 |
| BRCA1 Associated RING Domain 1 | BARD1 | Q99728 | 1095 | -1.13 |
| Cullin 3 | CUL3 | Q13618 | 345 | -1.13 |
| RNA binding motif protein 26 | RBM26 | Q5T8P6 | 897 | -1.14 |
| Bcl10-interacting CARD protein | C9ORF89 | Q96LW7 | 422 | -1.15 |
| S-phase kinase-associated protein 1 | SKP1A/p19 | P63208 | 363 | -1.16 |
| B melanoma antigen 3/ Cancer/testis antigen 2.3 | BAGE3 | Q86Y29 | 420 | -1.20 |
| Cytosolic Thiouridylase Subunit 1 | CTU1 | Q7Z7A3 | 907 | -1.20 |
| SUMO1-activating enzyme 2 | UBA2 | Q9UBT2 | 874 | -1.22 |
| Tissue inhibitor of metalloproteinases 2 | TIMP2 | P16035 | 537 | -1.22 |
| RING finger and WD repeat domain protein 2 | RFWD2 | Q8NHY2 | 1107 | -1.23 |
| Coagulation factor VII (light chain,Cleaved-Arg212) | FA7 | P08709 | 1156 | -1.29 |
| Rho guanine nucleotide exchange factor 3 | ARHGEF3 | Q9NR81 | 1123 | -1.30 |
| Cytochrome c oxidase assembly protein COX11 | COX11 | Q9Y6N1 | 911 | -1.31 |
| Diacylglycerol Kinase eta | DGKH | Q86XP1 | 908 | -1.36 |
| ATP-binding cassette sub-family A member 8 | ABCA8 | O94911 | 848 | -1.37 |
| Shugoshin-like 1 | SGOL1 | Q5FBB7 | 895 | -1.37 |
| Guanylate Cyclase Beta 1 | GUCY1B3 | Q02153 | 853 | -1.38 |
| Cell division cycle 7 | CDC7 | O00311 | 1044 | -1.38 |
| Aldo-keto reductase family 1 member C-like protein 1 | AKR1CL1 | Q5T2L2 | 246 | -1.41 |
| Interleukin 2 | IL2 | P60568 | 832 | -1.55 |
| Cytochrome P450 7B1 | CYP7B1 | O75881 | 802 | -1.82 |
| Ubiquitin-protein ligase E3B | UBE3B | Q7Z3V4 | 291 | -2.19 |

* The vertical lines demarcate the 1.15 fold change (FC) cut-off, resulting in 21 up-regulated and 19 down-regulated genes used in IPA

### Table S2: List of IPA predicted OFC-related genes

| Symbol | Entrez Gene Name | Location | Family | Entrez Gene ID for Human | Entrez Gene ID for Mouse |
|---|---|---|---|---|---|
| **ACVR2A** | activin A receptor, type IIA | Plasma Membrane | kinase | 92 | 11480 |
| **AHR** | aryl hydrocarbon receptor | Nucleus | ligand-dependent nuclear receptor | 196 | 11622 |
| **AKAP8** | A kinase (PRKA) anchor protein 8 | Nucleus | other | 10270 | 56399 |
| **ALX1** | ALX homeobox 1 | Nucleus | transcription regulator | 8092 | 216285 |
| **ALX4** | ALX homeobox 4 | Nucleus | transcription regulator | 60529 | 11695 |
| **Anp32b** | acidic (leucine-rich) nuclear phosphoprotein 32 family, member B | Nucleus | other | | 67628 |
| **ASPH** | aspartate beta-hydroxylase | Cytoplasm | enzyme | 444 | 65973 |
| **ASXL1** | additional sex combs like transcriptional regulator 1 | Nucleus | transcription regulator | 171023 | 228790 |
| **BARX1** | BARX homeobox 1 | Nucleus | transcription regulator | 56033 | 12022 |
| **BMP4** | bone morphogenetic protein 4 | Extracellular Space | growth factor | 652 | 12159 |
| **BMP7** | bone morphogenetic protein 7 | Extracellular Space | growth factor | 655 | 12162 |
| **CDH1** | cadherin 1, type 1, E-cadherin (epithelial) | Plasma Membrane | other | 999 | 12550 |
| **CDKN1A** | cyclin-dependent kinase inhibitor 1A (p21, Cip1) | Nucleus | kinase | 1026 | 12575 |
| **Cdkn1c** | cyclin-dependent kinase inhibitor 1C (P57) | Nucleus | other | | 12577 |
| **CHUK** | conserved helix-loop-helix ubiquitous kinase | Cytoplasm | kinase | 1147 | 12675 |
| **COL2A1** | collagen, type II, alpha 1 | Extracellular Space | other | 1280 | 12824 |
| **CRK** | v-crk avian sarcoma virus CT10 oncogene homolog | Cytoplasm | other | 1398 | 12928 |
| **CYP51A1** | cytochrome P450, family 51, subfamily A, polypeptide 1 | Cytoplasm | enzyme | 1595 | 13121 |
| **DHCR7** | 7-dehydrocholesterol reductase | Cytoplasm | enzyme | 1717 | 13360 |
| **DHFR** | dihydrofolate reductase | Nucleus | enzyme | 1719 | 13361 |
| **DLG1** | discs, large homolog 1 (Drosophila) | Plasma Membrane | kinase | 1739 | 13383 |
| **DLX1** | distal-less homeobox 1 | Nucleus | transcription regulator | 1745 | 13390 |
| **DNMT3B** | DNA (cytosine-5-)-methyltransferase 3 beta | Nucleus | enzyme | 1789 | 13436 |
| **DPH1** | diphthamide biosynthesis 1 | Cytoplasm | other | 1801 | 116905 |
| **EDN1** | endothelin 1 | Extracellular Space | cytokine | 1906 | 13614 |
| **EFNA5** | ephrin-A5 | Plasma Membrane | kinase | 1946 | 13640 |
| **EFNB1** | ephrin-B1 | Plasma Membrane | other | 1947 | 13641 |
| **EFTUD2** | elongation factor Tu GTP binding domain containing 2 | Nucleus | enzyme | 9343 | 20624 |
| **EGF** | epidermal growth factor | Extracellular Space | growth factor | 1950 | 13645 |
| **EGFR** | epidermal growth factor receptor | Plasma Membrane | kinase | 1956 | 13649 |
| **EPHB2** | EPH receptor B2 | Plasma Membrane | kinase | 2048 | 13844 |
| **EPHB3** | EPH receptor B3 | Plasma Membrane | kinase | 2049 | 13845 |
| **FGF10** | fibroblast growth factor 10 | Extracellular Space | growth factor | 2255 | 14165 |
| **FGF9** | fibroblast growth factor 9 | Extracellular Space | growth factor | 2254 | 14180 |
| **FGFR1** | fibroblast growth factor receptor 1 | Plasma Membrane | kinase | 2260 | 14182 |
| **FGFR2** | fibroblast growth factor receptor 2 | Plasma Membrane | kinase | 2263 | 14183 |
| **FIGN** | fidgetin | Nucleus | other | 55137 | 60344 |
| **folic acid** | | Other | chemical - endogenous mammalian | | |
| **FOXC2** | forkhead box C2 | Nucleus | transcription regulator | 2303 | 14234 |
| **FOXE1** | forkhead box E1 | Nucleus | transcription regulator | 2304 | 110805 |
| **GABRB3** | gamma-aminobutyric acid (GABA) A receptor, beta 3 | Plasma Membrane | ion channel | 2562 | 14402 |
| **GAD1** | glutamate decarboxylase 1 (brain, 67kDa) | Cytoplasm | enzyme | 2571 | 14415 |
| **GAS1** | growth arrest-specific 1 | Plasma Membrane | other | 2619 | 14451 |
| **GBX2** | gastrulation brain homeobox 2 | Nucleus | transcription regulator | 2637 | 14472 |
| **GDF11** | growth differentiation factor 11 | Extracellular Space | growth factor | 10220 | 14561 |
| **GLCE** | glucuronic acid epimerase | Cytoplasm | enzyme | 26035 | 93683 |
| **GRHL3** | grainyhead-like 3 (Drosophila) | Nucleus | other | 57822 | 230824 |
| **GSC** | goosecoid homeobox | Nucleus | transcription regulator | 145258 | 14836 |
| **GSK3B** | glycogen synthase kinase 3 beta | Nucleus | kinase | 2932 | 56637 |
| **H19** | H19, imprinted maternally expressed transcript (non-protein coding) | Cytoplasm | other | 283120 | 14955 |
| **HOXA2** | homeobox A2 | Nucleus | transcription regulator | 3199 | 15399 |
| **HSPG2** | heparan sulfate proteoglycan 2 | Extracellular Space | enzyme | 3339 | 15530 |
| **INPP5E** | inositol polyphosphate-5-phosphatase, 72 kDa | Cytoplasm | phosphatase | 56623 | 64436 |
| **INSIG1** | insulin induced gene 1 | Cytoplasm | other | 3638 | 231070 |
| **INSIG2** | insulin induced gene 2 | Cytoplasm | other | 51141 | 72999 |
| **IRF6** | interferon regulatory factor 6 | Nucleus | transcription regulator | 3664 | 54139 |
| **ITGB8** | integrin, beta 8 | Plasma Membrane | other | 3696 | 320910 |
| **JAG2** | jagged 2 | Extracellular Space | growth factor | 3714 | 16450 |
| **KAT6A** | K(lysine) acetyltransferase 6A | Nucleus | enzyme | 7994 | 244349 |
| **lovastatin** | | Other | chemical drug | | |
| **LRP6** | low density lipoprotein receptor-related protein 6 | Plasma Membrane | transmembrane receptor | 4040 | 16974 |
| **Map3k7** | mitogen-activated protein kinase kinase kinase 7 | Cytoplasm | kinase | | 26409 |
| **MDM2** | MDM2 proto-oncogene, E3 ubiquitin protein ligase | Nucleus | transcription regulator | 4193 | 17246 |
| **MDM4** | MDM4, p53 regulator | Nucleus | other | 4194 | 17248 |
| **MNT** | MAX network transcriptional repressor | Nucleus | transcription regulator | 4335 | 17428 |

| | | | | | |
|---|---|---|---|---|---|
| **MSTN** | myostatin | Extracellular Space | growth factor | 2660 | 17700 |
| **MSX1** | msh homeobox 1 | Nucleus | transcription regulator | 4487 | 17701 |
| **NABP2** | nucleic acid binding protein 2 | Nucleus | other | 79035 | 69917 |
| **NDST1** | N-deacetylase/N-sulfotransferase (heparan glucosaminyl) 1 | Cytoplasm | enzyme | 3340 | 15531 |
| **OSR2** | odd-skipped related transciption factor 2 | Nucleus | transcription regulator | 116039 | 107587 |
| **PBX1** | pre-B-cell leukemia homeobox 1 | Nucleus | transcription regulator | 5087 | 18514 |
| **PBX2** | pre-B-cell leukemia homeobox 2 | Nucleus | other | 5089 | 18515 |
| **PBX3** | pre-B-cell leukemia homeobox 3 | Nucleus | transcription regulator | 5090 | 18516 |
| **PDGFC** | platelet derived growth factor C | Extracellular Space | growth factor | 56034 | 54635 |
| **PDGFRA** | platelet-derived growth factor receptor, alpha polypeptide | Plasma Membrane | kinase | 5156 | 18595 |
| **PHC1** | polyhomeotic homolog 1 (Drosophila) | Nucleus | transcription regulator | 1911 | 13619 |
| **PITX2** | paired-like homeodomain 2 | Nucleus | transcription regulator | 5308 | 18741 |
| **RAD23B** | RAD23 homolog B (S. cerevisiae) | Nucleus | other | 5887 | 19359 |
| **RAX** | retina and anterior neural fold homeobox | Nucleus | transcription regulator | 30062 | 19434 |
| **RECQL4** | RecQ protein-like 4 | Nucleus | enzyme | 9401 | 79456 |
| **retinoid** | | Other | chemical drug | | |
| **RPGRIP1L** | RPGRIP1-like | Cytoplasm | other | 23322 | 244585 |
| **RSPO2** | R-spondin 2 | Extracellular Space | other | 340419 | 239405 |
| **RYK** | receptor-like tyrosine kinase | Plasma Membrane | kinase | 6259 | 20187 |
| **SATB2** | SATB homeobox 2 | Nucleus | transcription regulator | 23314 | 212712 |
| **SC5D** | sterol-C5-desaturase | Cytoplasm | enzyme | 6309 | 235293 |
| **SHH** | sonic hedgehog | Extracellular Space | peptidase | 6469 | 20423 |
| **SLC32A1** | solute carrier family 32 (GABA vesicular transporter), member 1 | Plasma Membrane | transporter | 140679 | 22348 |
| **SNAI1** | snail family zinc finger 1 | Nucleus | transcription regulator | 6615 | 20613 |
| **SNAI2** | snail family zinc finger 2 | Nucleus | transcription regulator | 6591 | 20583 |
| **SOX11** | SRY (sex determining region Y)-box 11 | Nucleus | transcription regulator | 6664 | 20666 |
| **SUMO1** | small ubiquitin-like modifier 1 | Nucleus | enzyme | 7341 | 22218 |
| **TBX1** | T-box 1 | Nucleus | transcription regulator | 6899 | 21380 |
| **TBX10** | T-box 10 | Nucleus | transcription regulator | 347853 | 109575 |
| **TBX22** | T-box 22 | Nucleus | transcription regulator | 50945 | 245572 |
| **TCOF1** | Treacher Collins-Franceschetti syndrome 1 | Nucleus | transporter | 6949 | 21453 |
| **TCTN2** | tectonic family member 2 | Extracellular Space | other | 79867 | 67978 |
| **TGFA** | transforming growth factor, alpha | Extracellular Space | growth factor | 7039 | 21802 |
| **TGFB1** | transforming growth factor, beta 1 | Extracellular Space | growth factor | 7040 | 21803 |
| **TGFB3** | transforming growth factor, beta 3 | Extracellular Space | growth factor | 7043 | 21809 |
| **TGFBR2** | transforming growth factor, beta receptor II (70/80kDa) | Plasma Membrane | kinase | 7048 | 21813 |
| **TP63** | tumor protein p63 | Nucleus | transcription regulator | 8626 | 22061 |
| **UBB** | ubiquitin B | Cytoplasm | enzyme | 7314 | |
| **VAX1** | ventral anterior homeobox 1 | Nucleus | transcription regulator | 11023 | 22326 |
| **VAX2** | ventral anterior homeobox 2 | Nucleus | transcription regulator | 25806 | 24113 |
| **VEGFA** | vascular endothelial growth factor A | Extracellular Space | growth factor | 7422 | 22339 |
| **WDPCP** | WD repeat containing planar cell polarity effector | Plasma Membrane | other | 51057 | 216560 |
| **WFIKKN1** | WAP, follistatin/kazal, immunoglobulin, kunitz and netrin domain containing 1 | Cytoplasm | other | 117166 | 215001 |
| **WFIKKN2** | WAP, follistatin/kazal, immunoglobulin, kunitz and netrin domain containing 2 | Extracellular Space | other | 124857 | 278507 |
| **WNT9B** | wingless-type MMTV integration site family, member 9B | Extracellular Space | other | 7484 | 22412 |

Table S3: IPA predicted diseases and bio-functions associated with the 40 gene-products significantly altered in Topiramate-treated HEPM cells.

**DISEASES AND BIO FUNCTIONS**

| Rank (IPA) | Categories | Functions | Diseases or Functions Annotation | p-Value | Predicted Activation State | Activation z-score | Bias-corrected z-score | Molecules | No. of Molecules |
|---|---|---|---|---|---|---|---|---|---|
| 1 | Cellular Development, Hematological System Development and Function, Hematopoiesis | maturation | maturation of effector lymphocytes | 4.96E-08 | | | | IRF4, TGFB1, IL2 | 3 |
| 6 | Organismal Survival | lifespan | lifespan of organism | 7.22E-06 | | -1.205 | -1.241 | TGFB1, IL2, SOD1, RET, GUCY1B3 | 5 |
| 7 | Cell Death and Survival, Nervous System Development and Function | cell viability | cell viability of neurons | 7.34E-06 | | 0.658 | 0.591 | TGFB1, IL2, PTCH1, SOD1, RET, CHEK2 | 6 |
| 13 | Cell Death and Survival | survival | cell survival | 1.13E-05 | | -0.01 | -0.106 | IRF4, TYRO3, PTCH1, CDC7, F7, SOD1, TRADD, IL2, TGFB1, TNFSF8, RET, CHEK2, TIMP2 | 13 |
| 29 | Cell Death and Survival | cell viability | cell viability | 0.0000303 | | -0.035 | -0.132 | TRADD, IRF4, TGFB1, IL2, TNFSF8, PTCH1, CDC7, F7, SOD1, RET, CHEK2, TIMP2 | 12 |
| 37 | Cell Death and Survival | apoptosis | apoptosis | 0.0000569 | | 0.122 | 0.15 | IRF4, TYRO3, PTCH1, CDC7, TNNI3, F7, SOD1, TRADD, GADD45GIP1, IL2, TGFB1, TNFSF8, RFWD2, ARHGEF3, RET, CHEK2, TIMP2 | 17 |

Table S4: IPA predicted networks associated with the 40 gene-products significantly altered in Topiramate-treated HEPM cells.

| Networks ID (IPA) | Molecules in Network | Score | Focus Molecules | Top Diseases and Functions |
|---|---|---|---|---|
| 1 | ADRB, ANAPC1, ARHGEF3, Akt, CD3, CDC7, CDH10, Cdc2, Creb, Cyclin A, Cyclin B, GADD45GIP1, GUCY1B3, Histone h3, Histone h4, Hsp70, Hsp90, NFkB (complex), PI3K (complex), PTCH1, RET, RFWD2, RNA polymerase II, SGOL1, SKP1, SOD1, SRC (family), TCR, TRADD, UBA2, Ubiquitin, Vegf, caspase, estrogen receptor, p85 (pik3r) | 32 | 14 | Cancer, Endocrine System Disorders, Organismal Injury and Abnormalities |
| 2 | Ap1, BCR (complex), CYP7B1, Calcineurin protein(s), Collagen type I, Collagen type IV, Collagen(s), Cyclin E, ERK1/2, F7, IFN Beta, IL1, IL12 (complex), IL12 (family), IL2, IL23, IRF4, IgG, Iga, Ige, Igm, Immunoglobulin, Interferon alpha, LDL, MHC Class II (complex), NOX5, Nfat (family), P38 MAPK, PDGF BB, TIMP2, TNFSF8, TYRO3, Tgf beta, Tnf (family), Tnf receptor | 16 | 8 | Cell Death and Survival, Cellular Development, Hematological System Development and Function |
| 3 | ABCF1, ADH7, ALDH3B1, APP, ATP5D, CAND1, CELF2, CSRP1, CTU1, DPM2, DUSP3, EED, EXOSC3, FBXL20, GTPBP1, HIVEP3, ITM2B, MYO1D, NIFK, Nrgn, ORAOV1, PDCD11, PIGH, PNO1, POLH, PSAT1, REG1A, Rplp1 (includes others), SLC11A2, SLC13A3, SLX4, UBC, VKORC1, YAE1D1, alcohol dehydrogenase | 13 | 7 | Cell Morphology, Cellular Assembly and Organization, Neurological Disease |
| 4 | ADCY9, AKAP10, ARHGAP1, ATP9A, BCL10, C9orf89, CHEK2, CHST8, COX11, DGKH, DUSP7, ERK, FILIP1L, FSH, GPRC5B, HYAL2, Insulin, Jnk, LRRC32, Lh, MSMB, Mapk, PPP2R5B, Pka, Pkc(s), Ras, SLC52A2, SLC5A2, STK17A, TGFB1, TNNI3, VTCN1, XPNPEP2, cytochrome-c oxidase, hemoglobin | 11 | 6 | Cell Cycle, Hair and Skin Development and Function, Carbohydrate Metabolism |
